# Mass spectrometric detection of protein nanoclusters: the invisible intermediates of phase separation

**DOI:** 10.64898/2026.10.28.762497

**Authors:** Christina Glen Robb, Anisha Haris, Thuy P Dao, Carlos Castañeda, Jakub Ujma, Rebecca Beveridge

## Abstract

Phase separation of biomolecules, including intrinsically disordered proteins, is an underlying physical process behind cellular condensate formation. To increase mechanistic understanding of its initial stages, we demonstrate the application of charge detection mass spectrometry to characterise the mass dispersion of protein nanoclusters that represent intermediates in this important biophysical process. We detect polydisperse clusters of 1–280 protein molecules (64 kDa–18.4 MDa), that are currently invisible to most analytical techniques.

## Main text

Phase separation is a vital and ubiquitous biological process in which a homogenous solution containing macromolecules (i.e. proteins, RNA, DNA) separates into distinct macromolecule-dilute and macromolecule-dense phases. These macromolecule-dense phases (with material properties ranging from liquid-like to solid) can form biomolecular condensates in cells.^1,2^ Phase separation regulates essential biological processes and aberrant phase separation is associated with many pathologies.^3,4^ Despite the biological importance of phase separation, mechanistic understanding of its initial stages remains elusive due to challenges in analysing soluble protein species at sub-saturating concentrations. This is partially because initially formed protein nanoclusters, which cannot be detected with light microscopy, are highly polydisperse and hence evade characterisation with most analytical methods.^5^ Additionally, many phase separating biomolecules are highly dynamic intrinsically disordered proteins (IDPs) and therefore their interactions are challenging to characterise with the current range of biophysical tools.^6^

Protein nanoclusters that form below the protein concentration required for phase separation have been independently identified in multiple systems, including the FET family of proteins^7^ and α-synuclein.^8^ Current tools for nanocluster analysis *in vitro* include dynamic light scattering (DLS), mass photometry and microfluidic resistive pulse sensing (MRPS). The diffusion speed of nanoclusters measured by DLS can be converted into a hydrodynamic size and, through the use of certain density approximations, provides a distribution of the number of protein molecules per cluster.^9^ This information has been useful for comparing cluster sizes in solutions of different cations.^5,10^ Mass photometry provides a more direct readout of nanocluster size based on the measured mass distribution. However, a caveat here is that proteins with monomeric M_w_ above the lower limit of the mass photometer (40 kDa) saturate the detector, rendering analysis infeasible. If the monomer is below 40 kDa, as is the case for α-synuclein, mass photometry enables efficient nanocluster characterisation but cannot report on the lower-mass species.^8^ MRPS quantifies particle size distributions by measuring the change in electrical resistance as particles travel through a nano-constriction, allowing quantification of sizes from 10 nm to 10 µm. This has been used to quantify nanoclusters of MTRN3, nuclear speckle proteins and synapsin.^11–13^

Here we demonstrate that charge detection mass spectrometry (CDMS)^14^ is uniquely suited to quantitatively characterise the extreme polydispersity of phase separating protein systems, by simultaneously reporting on the precise stoichiometry of lower-order oligomers of n=1–6 and the distribution of higher-order protein nanoclusters up to n=280.^15^ Such analysis is enabled by CDMS simultaneously measuring the mass to charge ratio (*m*/*z*) and the charge (*z*) of individual ions, thus providing direct mass measurements of individual protein complexes and clusters.^16,17^ CDMS has been used to study virus like-nanoparticles, adeno-associated virus capsids and protein aggregates.^15,18–20^ Here we use CDMS to measure complexes and clusters formed by ubiquilin-2 (UBQLN2), which represents an ideal model protein for our studies. UBQLN2 contains extensive regions of intrinsic disorder and its phase separation, which is promoted by high ionic strength solutions, is well-characterised *in vitro* and in cells.^21–23^

To test the capability of CDMS for characterising protein nanoclusters, we analysed UBQLN2 from a low-salt solution (10 mM ammonium acetate, AmAc) at which it’s expected to remain fully soluble (Figure 1a inset), and high-salt (75 mM AmAc) at which small puncta can be seen by bright field microscopy (Figure 1b inset) but macroscopic phase separation does not occur. At low salt (Figure 1a), a narrow distribution of masses and charges is detected; of the 4066 ions, 75% fall below 150 kDa, corresponding to the UBQLN2 dimer.^22,24^ Whilst a similar number of ions (3762) were detected upon phase separation induction (high-salt) (Figure 1b), only 10% of the data points fall below 150 kDa, and a shift to a wider range of charges and masses is apparent. The largest ion at low-salt is 5.98 MDa, (91 UBQLN2 molecules), increasing to 18.42 MDa, (280 UBQLN2 molecules) at high-salt. This shift in distribution is also seen in the mass histograms (Figure 1c), which show that the median mass of UBQLN2 complexes is 131 kDa at low-salt (blue histogram), and 882 kDa at high-salt (orange histogram) with the latter corresponding to a mass of ∼14 UBQLN2 molecules.

**Figure 1:**
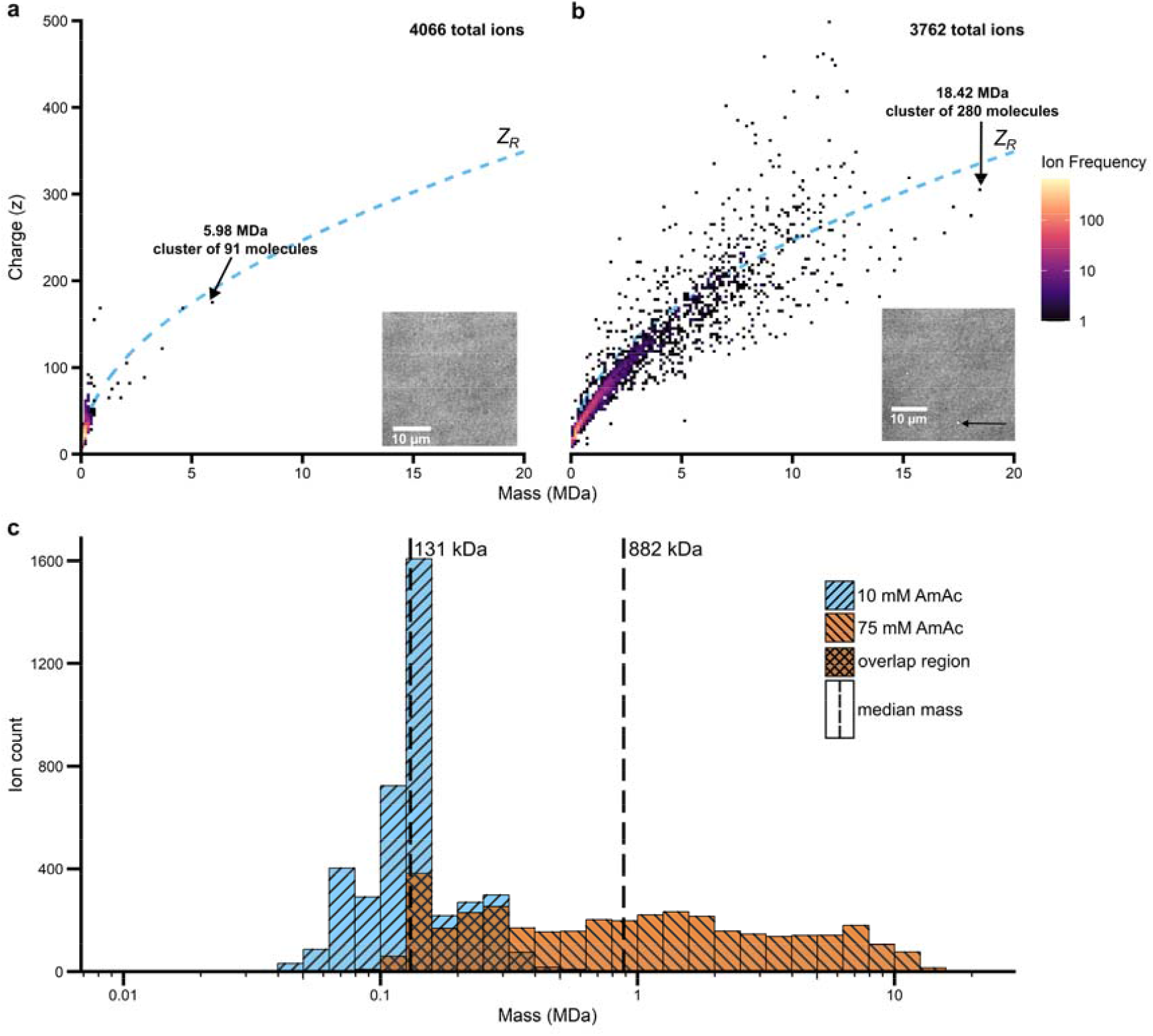
Charge-to-mass heat maps of UBQLN2 (15 µM) from a) 10 mM and b) 75 mM AmAc. The largest ions detected are identified by an arrow, accompanied with their mass in MDa and the number of UBQLN2 molecules in the cluster. Insets: Microscopy images of UBQLN2 in equivalent solutions, with puncta indicated by an arrow (taken from reference 22 under creative commons licence CC BY 4.0). The dashed line in a and b corresponds to the Rayleigh limit (Z_R_). c) Mass histograms of UBQLN2 ions from 10 mM (blue) and 75 mM (orange) AmAc. Ion counts are shown in logarithmically spaced mass bins (0.1 log_10_ units wide) on a log_10_ mass axis. Dashed vertical lines indicate the median mass of each data set (131 kDa and 882 kDa for 10 mM and 75 mM, respectively). Overlapping regions of the two distributions appear darker and with a checked pattern.

Whilst the mass dispersion of the nanoclusters is highly informative, their charge distribution adds a further dimension to the analysis in terms of their overall shape. The Rayleigh limit (*Z*_*R*_) is the maximum number of charges that a spherical ion can accommodate, meaning that protein ions with a charge state below the *Z*_*R*_ are likely spherical and compact, and those with a charge state above the *Z*_*R*_ are likely elongated.^25^ Therefore, the charge distribution of the nanoclusters formed in 75 mM

AmAc is intriguing. Up to a mass of ∼5 MDa, there is a preference of the nanoclusters to have a charge below *Z*_*R*_, indicating that these clusters are compact. Above 5 MDa, there is a broad charge distribution across the mass range, with data points quite evenly distributed below and above *Z*_*R*_. This indicates that larger nanoclusters have broader ranges of geometrical dimensions than their lower-mass counterparts. Whilst highly-charged species of thermally-induced bovine serum albumin (BSA) aggregates (incubated at 5 mg/ml at 75 °C for 20 minutes) have also been identified by Jordan et al., the majority of MDa-scale analytes have charge states close to or lower than the *Z*_*R*_, suggesting compact conformations.^17,20,26^

To test the generality of nanocluster formation we also measured BSA; a globular protein of equivalent mass to UBQLN2 that doesn’t readily undergo phase separation (Figure S1). BSA forms oligomers up to a tetramer, with the pentamer and hexamer also present in <10 data points. In comparison, UBQLN2 forms larger oligomers even in non-phase separating conditions, indicating that UBQLN2 and possibly other phase separating IDPs form nanoclusters in the solutions currently considered to confer full solubility.

Next, we were interested to investigate the ability of CDMS to detect the lower-order oligomers that may represent the building blocks for the larger nanoclusters. In the solution that favours full protein solubility (low salt, Figure 2a) the mass distribution profile shows oligomers from monomer to hexamer, with complexes corresponding to an even number of proteins favoured but odd numbers also present. When the salt concentration is increased to induce phase separation (Figure 2b), there is a decrease in the pentamer and hexamer abundance across both measurements, suggesting their propensity to progress to larger nanoclusters.

**Figure 2:**
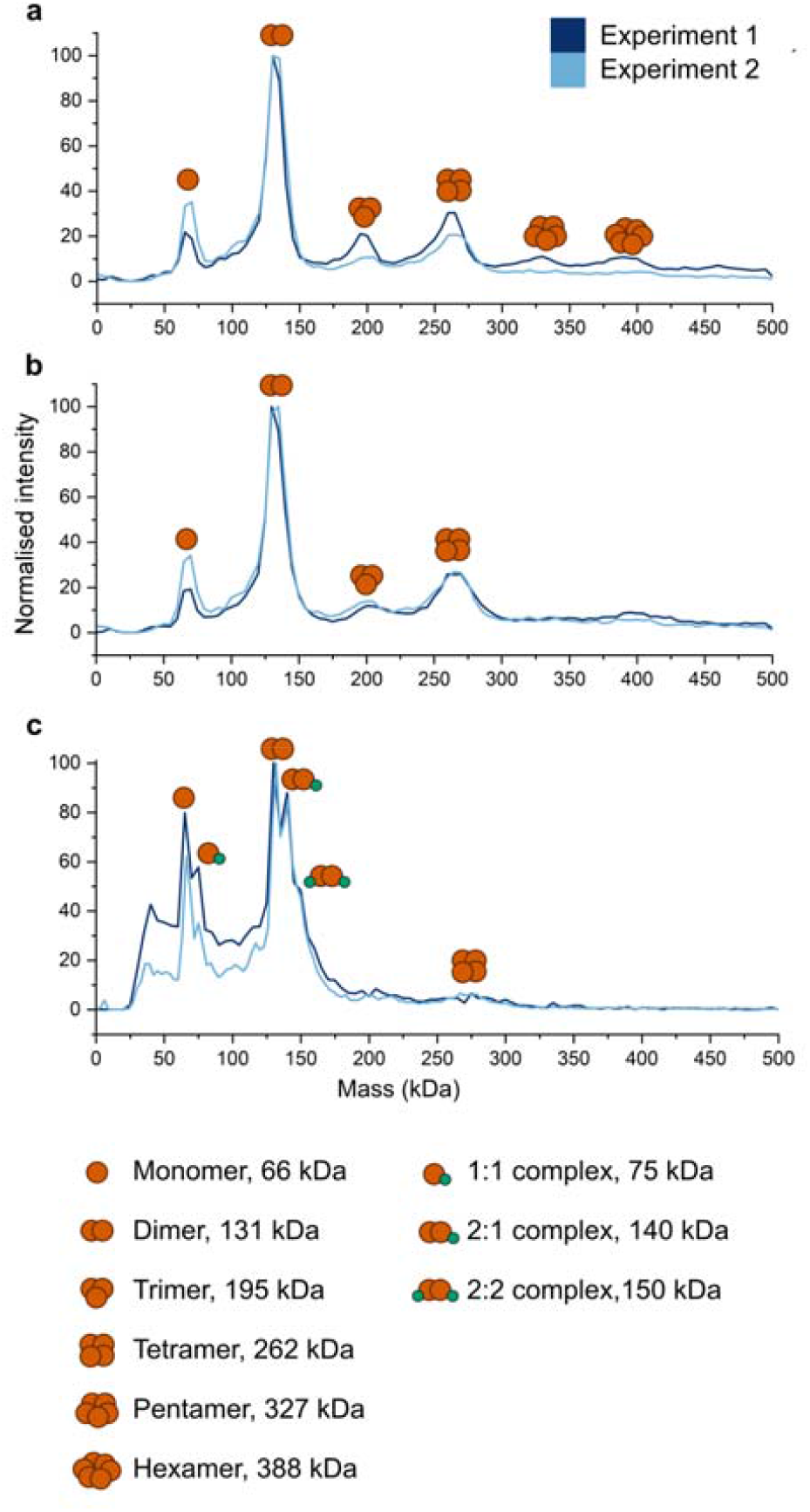
Mass distribution profiles of UBQLN2 from a) 10 mM AmAc, b) 75 mM AmAc. c) 1:4 UBQLN2:Ub from 10 mM AmAc. Results from two repeats (n = 2) are shown in dark and light blue. Peaks are annotated with the assigned oligomer or complex.

Upon addition of Ub (Figure 2c), which is a potent inhibitor of UBQLN2 phase separation,^23^ signal corresponding to 1:1, 2:1 and 2:2 UBQLN2 to Ub stoichiometries is observed.

Additionally, in comparison to UBQLN2 alone, no signal is present corresponding to the UBQLN2 trimer, pentamer and hexamer and signal corresponding to the UBQLN2 tetramer is significantly depleted. This allows us to deduce that Ub binding favours the monomer and dimer of UBQLN2 and disrupts higher-order oligomers. The mass vs charge scatter plot for UBQLN2 + Ub is similar to that of UBQLN2 alone in 10 mM AmAc (Figure S2).

The results corresponding to the UBQLN2:Ub complexes are highly complementary to data from previous MS experiments in which equivalent complexes were reported.

However, whilst we were confident in assigning the 2:1 UBQLN2 to Ub complexes, the low abundance of the 2:2 and 1:1 complexes along with their coincident *m/z* assignments rendered their detection to be reported with lower confidence.^22^ This highlights the strength of CDMS in elucidating multiple complexes present in a stoichiometric mixture.

With CDMS we have quantified the mass dispersion of protein nanoclusters that represent intermediate species in phase separation. The ion distribution for the nanoclusters is stochastic and indicates that UBQLN2 nanoclusters do not form in a uniform manner, and rather that the number of protein molecules present in the cluster can vary widely. CDMS will be highly informative for analysing additional phase separating proteins including those with pathological mutations and those involved in designer or smart condensates.

## Copyright Statement and Competing Interests

XEVO is a trademark of Waters Corporation or its affiliates. JU and AH are employees of Waters Corporation.

## Acknowledgements

R.B. is grateful the receipt of a Phillip Leverhulme Prize from the Leverhulme Trust, which funded this project. R.B. is also supported by a UKRI Future Leaders Fellowship (MR/T020970/1) and its renewal (MR/Y034236/1). C.A.C. acknowledges support from National Institutes of Health (NIH) R35 GM158070. The content is solely the responsibility of the authors and does not necessarily represent the official views of the NIH.

