## Supplementary Information for "Mass spectrometric detection of protein nanoclusters: the invisible intermediates of phase separation"

a. Department of Pure and Applied Chemistry, University of Strathclyde, Glasgow G1 1XL, U.K.
b. Waters Corporation, Stamford Avenue, Altrincham Road, Wilmslow SK9 4AX, U.K.
c. Departments of Biology and Chemistry, BioInspired Institute, Syracuse University, Syracuse, New York 13244, United States.

The supporting information includes:

Methods and Materials

Supplementary Figure S1 to S3

**Methods**

**Protein expression, and purification**

Wildtype Ub was expressed and purified as detailed elsewhere.^1^ Full-length UBQLN2 was expressed and purified as described previously.^2^  Briefly, the construct was expressed in *E. coli* Rosetta 2 (DE3) pLysS cells in LB broth at 37 °C overnight. Bacteria were pelleted, frozen, lysed, then purified *via* a “salting out” process. NaCl was added to the cleared lysate to the final concentration of 0.5 M–1 M. UBQLN2 droplets were pelleted and then resuspended in 20 mM NaPhosphate, 0.5 mM EDTA, 0.1 mM TCEP, 0.02% NaN_3_ (pH 6.8). Leftover NaCl was removed through HiTrap desalting column (GE Healthcare). Purified proteins were frozen at −80 °C.

**Preparation of proteins for CDMS**

UBQLN2 (45 µM) was buffer exchanged into 10 mM AmAc pH 6.8 using 96-well micro dialysis plates (Thermo Fisher Scientific, Waltham, MA USA). Ub (100 µM) was buffer exchanged using Bio-Rad Micro Bio-Spin P6 columns (Bio-Rad, Hercules, CA, USA). Final protein concentrations were determined using a Implen NP80 nanophotometer (Implen, München, Germany) using the A280 method. Protein concentrations were calculated with extinction coefficients of 11460 M^−1^ cm^−1^ for UBQLN2 and 1490 M^−1^ cm^−1^ for Ub.

UBQLN2 was diluted to a protein concentration of 15 µM and final AmAc concentrations of 10 mM and 75 mM (pH 6.8) for experiments on the unbound protein. For experiments with UBQLN2 and Ub, a 1:4 molar ratio of UBQLN2 monomer to Ub was mixed by adding 15 µM of UBQLN2 (10 mM AmAc) to an equal volume of 60 µM of Ub (10 mM AmAc). Samples were allowed to equilibrate on ice for at least 30 minutes. Lyophilised bovine serum albumin (BSA) (Merck, P/N: 05470) was dissolved in 75 mM AmAc solution to a final protein concentration of 15 μM.

**CDMS Measurements**

CDMS data were acquired on a Waters Xevo^TM^ CDMS Instrument with an electrostatic linear ion trap (ELIT) (Waters Corporation, Wilmslow, UK).^3^ Nanoelectrospray emitters were pulled in-house with a Flaming/Brown P-97 micropipette puller from thin-walled glass capillaries (i.d. 0.78 mm, o.d. 1.0 mm, 10 cm length, both from Sutter Instrument Co., Novato, CA, USA). 10 µL of each sample was aspirated by a 25 µL gas-tight syringe (Hamilton^TM^, p/n 80265) and injected into the emitter via the syringe port of the nESI source. Wash solutions were (A) deionised water, (B) magic mix (1:1:1:1 mixture of water : isopropanol : methanol : acetonitrile), and (C) 10 or 75 mM AmAc, depending on the sample solution. The wash sequence (A, B, A, C) was performed between each sample injection, using 250 µL syringe (Hamilton, p/n 81165). A positive potential of 1.2–1.7 kV was applied to the solution via an internal stainless-steel capillary. Ions were trapped in the ELIT for 100 ms (Figure 1, 2a and 2b) or 400 ms (Figure 2c), where longer trapping time results in increased mass resolution, allowing ions with similar masses to be distinguishable. The frequency and amplitude information were converted to *m/z* and *z* values, respectively, and ultimately mass (*m/z* × *z*). These data were subsequently binned to generate the corresponding spectra (histograms). Spectra were collected until a minimum of approximately 3000 ions were recorded in total.

**Data processing**

Signal processing and data visualization were performed using waters_connect™ CDMS data viewer software. For the mass histograms, bin width was set to 5 kDa for UBQLN2 alone (Figures 2a and 2b) and 3 kDa for UBQLN2:Ub complex (Figure 2c). Maximum charge standard deviation was set to 4 for all data sets. Scatter plots in the extended data and mass histograms in Figures 2a, 2b and 2c were extracted from waters_connect™ and plots were created in OriginPro 2022 (OriginLab Corporation, Northampton, MA, USA). A custom R script^4^ in RStudio (RStudio Team, Boston, MA)^5^ was used to produce the heat maps and histogram plots in Figure 1. Final graphical editing and labelling was performed in Inkscape version 1.3.2 (Inkscape, [https://inkscape.org/](https://inkscape.org)).

**Supplementary Figures**

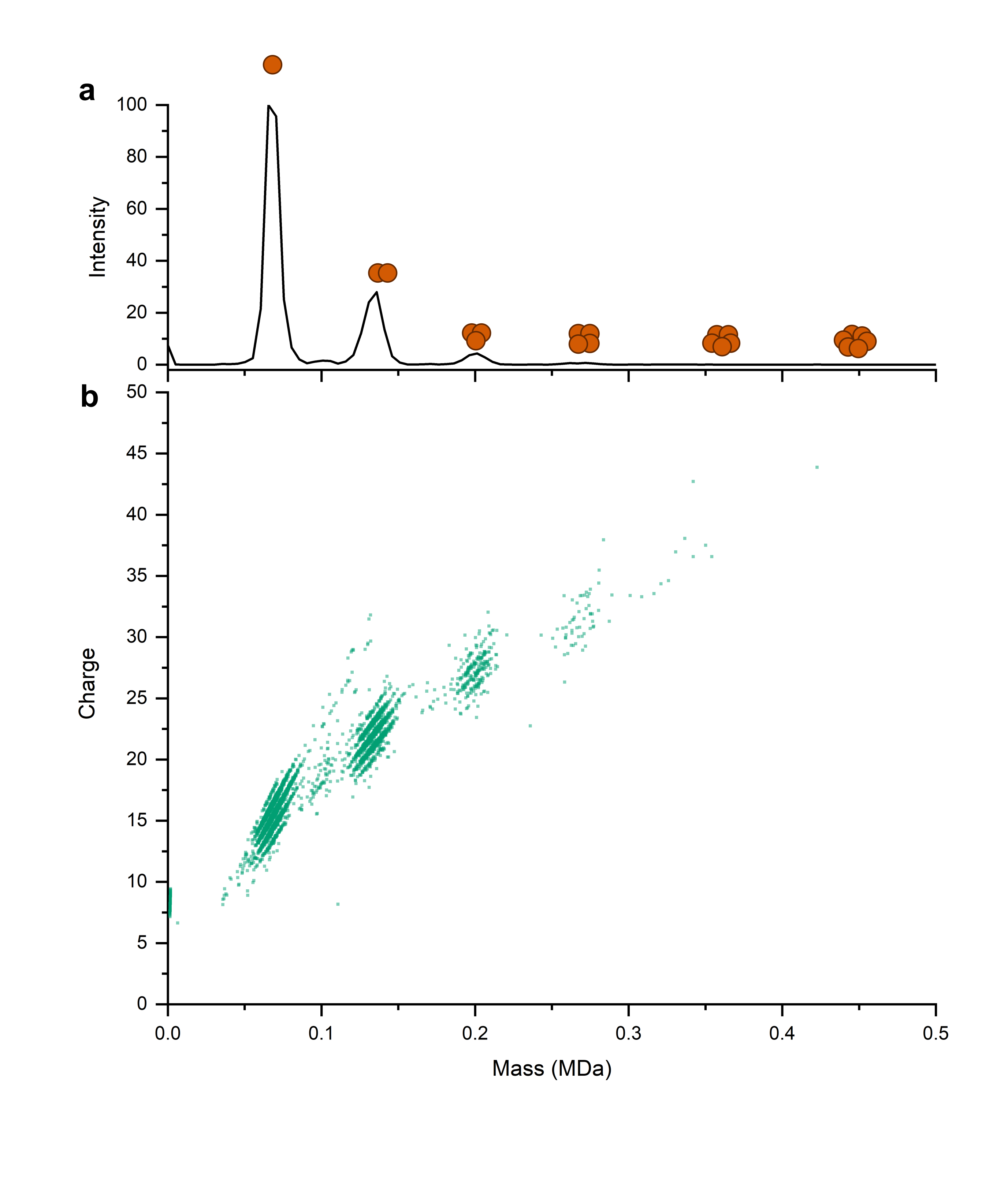

Figure S1 CDMS mass histogram (a) and mass vs charge scatter plot with 50% plot transparency (b) of 15 µM BSA from 75 mM AmAc. Monomer to tetramer is observed in high abundance and the pentamer and hexamer is observed in single digit ions. Almost all ions are below the Z_R_.

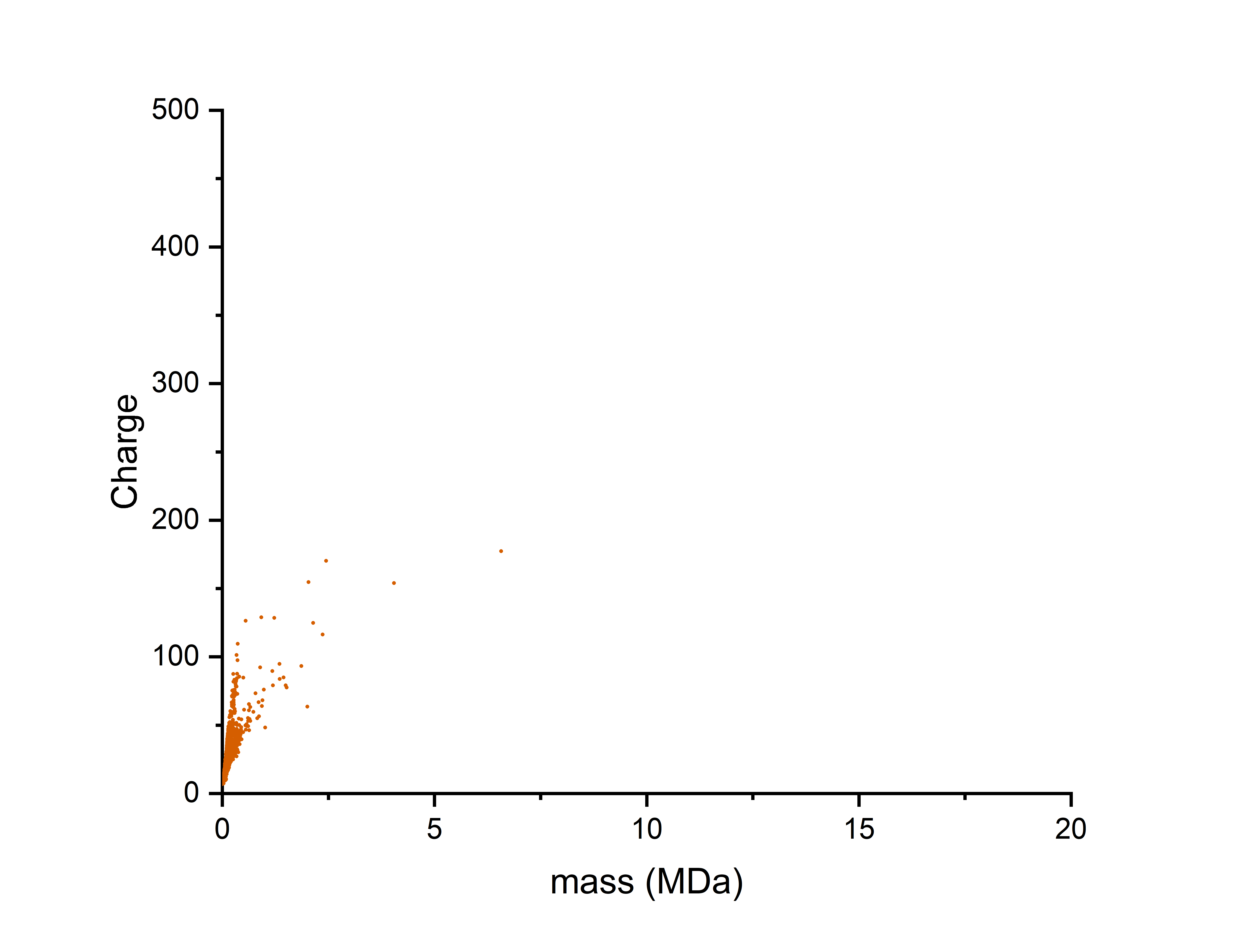

Figure S2 CDMS scatter plot of 1:4 molar ratio UBQLN2:Ub from 10 mM AmAc. Mass and charge density is comparable to that of UBQLN2 in 10 mM AmAc alone.

(1)	Castañeda, C. A.; Dixon, E. K.; Walker, O.; Chaturvedi, A.; Nakasone, M. A.; Curtis, J. E.; Reed, M. R.; Krueger, S.; Cropp, T. A.; Fushman, D. Linkage via K27 Bestows Ubiquitin Chains with Unique Properties among Polyubiquitins. *Structure* **2016**, *24* (3), 423–436. https://doi.org/10.1016/j.str.2016.01.007

(2)	Dao, T. P.; Kolaitis, R. M.; Kim, H. J.; O’Donovan, K.; Martyniak, B.; Colicino, E.; Hehnly, H.; Taylor, J. P.; Castañeda, C. A. Ubiquitin Modulates Liquid-Liquid Phase Separation of UBQLN2 via Disruption of Multivalent Interactions. *Molecular Cell* **2018**, *69* (6), 965–978. https://doi.org/10.1016/j.molcel.2018.02.004

(3)	Ujma, J.; Wheeldon, C.; Schofield, A.; Danby, M.; Eatough, D.; Bruton, D.; Haris, A.; Richardson, K.; Langridge, D.; Jarrell, A.; Brown, J. M.; Draper, B. E.; Jarrold, M. F.; Giles, K. A Charge Detection Mass Spectrometer for the Analysis of Megadalton-Sized Molecules. Biophysics June 1, 2026. https://doi.org/10.64898/2026.05.28.728353

(4)	R Core Team. *R: A Language and Environment for Statistical Computing.*; R Foundation for Statistical Computing: Vienna, Austria, 2025.

(5)	RStudio Team. *RStudio: Integrated Development for R.*; RStudio, Inc.: Boston, MA, 2019.
